# Extracting biological age from biomedical data via deep learning: too much of a good thing?

**DOI:** 10.1101/219162

**Authors:** Tim Pyrkov, Konstantin Slipensky, Mikhail Barg, Alexey Kondrashin, Boris Zhurov, Alexander Zenin, Mikhail Pyatnitskiy, Leonid Menshikov, Sergei Markov, Peter O. Fedichev

## Abstract

Aging-related physiological changes are systemic and, at least in humans, are linearly associated with age. Therefore, linear combinations of physiological measures trained to estimate chronological age have recently emerged as a practical way to quantify aging in the form of biological age. Aging acceleration, defined as the difference between the predicted and chronological age was found to be elevated in patients with major diseases and is predictive of mortality. In this work, we compare three increasingly accurate biological age models: metrics derived from unsupervised Principal Components Analysis (PCA), alongside two supervised biological age models; a multivariate linear regression and a state-of-the-art deep convolution neural network (CNN). All predictions were made using one-week long locomotor activity records from a 2003-2006 National Health and Nutrition Examination Survey (NHANES) dataset. We found that application of the supervised approaches improves the accuracy of the chronological age estimation at the expense of a loss of the association between the aging acceleration predicted by the model and all-cause mortality. Instead, we turned to the NHANES death register and introduced a novel way to train parametric proportional hazards models in a form suitable for out-of-the-box implementation with any modern machine learning software. Finally, we characterized a proof-of-concept example, a separate deep CNN trained to predict mortality risks that outperformed any of the biological age or simple linear proportional hazards models. Our findings demonstrate the emerging potential of combined wearable sensors and deep learning technologies for applications involving continuous health risk monitoring and real-time feedback to patients and care providers.

## INTRODUCTION

Many physiological parameters demonstrate profound correlations with age. This observation has led to the growing popularity of “biological clocks”, designed as linear predictors of chronological age from physiologically relevant variables, see, e.g., DNA methylation [1], gene expression profiles [2], plasma proteome [3]. However, a broader acceptance of the technology will depend on a better understanding of the observed correlations to the incidence of specific diseases, improved transferability of the models across populations and reduction of costs of the studies. We note, that large-scale biochemical or genomic profiling is not impossible, but is still logistically difficult and expensive. Instead, the recent introduction of low-power and compact sensors, based on micro-electromechanical systems (MEMS) has led to a new breed of the wearable and affordable devices providing unparalleled opportunities for the collecting and cloud-storing personal digitized activity records in a fully standardized and controlled way. This tracking is already done without interfering with the daily routines of hundreds of millions of people all over the world. Moreover, the analysis of human locomotion has already led to widely accepted health recommendations, such as the “10000 steps per day” minimum activity advisory [4], which is now a basic health recommendation in many wellness applications. Further, we have recently shown that a simple set of hand-engineered features, representing statistical properties of one-week long human physical activity time series can be used to produce digital biomarkers of aging and frailty [5].

Deep learning is a powerful tool in pattern recognition and has demonstrated outstanding performance in visual object identification, speech recognition, and other fields requiring hierarchical analysis of input data. Recent promising examples in the field of biomedical signal analysis include convolution neural networks (CNNs) trained to process electrocardiograms showing cardiologist-level performance in detection of arrhythmia [6], biomarkers of age from clinical blood biochemistry [7, 8] or electronic medical records [9], or mortality prediction [10]. Inspired by these examples, we explored deep learning architectures for Health Risks Assessment (HRA) applications, to use human physical activity streams from wearable devices for continuous health risks evaluation. First, we trained a series of deep CNNs to predict the chronological age of NHANES participants and obtained a substantial improvement over a multivariate linear regression fit from the same signal. By design, every such supervised biological age estimation aims at minimizing the difference between the predicted and the actual chronological age of the same patient. This quantity is referred to as aging acceleration and is associated with prevalence of disease [11–14] and mortality [15–17]. Therefore, we found, as expected, that any improvement yielding a decrease in the age prediction error rate often translates into a reduced ability to differentiate among disease states or mortality risks.

Thus we conclude that the chronological age of a patient is not the best target for biological age model development. More sensible ways to quantify aging involve the inclusion of metrics characterizing life expectancy, disease burden or frailty. Thus, we focused on mortality as the ultimate health status variable and proposed a novel technique to train a proportional hazards model in a form, suitable for an out-of-the-box implementation with modern deep-learning toolkits. In particular, we used the death register and raw one-week long time series representing physical activity records of NHANES patients and obtained the best results with a deep CNN architecture, simultaneously and automatically producing the set of features engineered from the raw physical activity time series and the most accurate non-linear representation of the risks function. Our findings demonstrate the emerging potential of a combination of wearable sensors and deep learning technologies for future HRA applications involving continuous health risk evaluation and real-time feedback to patients and care providers.

## RESULTS

### Deep learning chronological age from physical activity records

The “biological age” or “bioage” is a quantitative measure of aging and thus an expected lifespan based on biological data. State-of-the-art approaches for biological age evaluation take advantage of the strong association of physiological variables with age and thus rely on linear (see, e.g., DNA methylation age [1, 18]) or non-linear regressions [7, 8, 10] to estimate the chronological age of a patient directly from the biological data. Following these examples, we started by building a deep convolution neural network (CNN) trained to predict the chronological age of the same NHANES participants from the raw one-week long physical activity records. We relied on the CNN to automatically extract relevant features from time series (see, e.g., [6]) and to unravel the apparent non-linear dependencies between the locomotor activity variables and age. We trained a CNN_Age with four convolution and two dense layers (see Materials and Methods for the details of the CNN architecture). The model achieved Pearson’s *r* = 0.75 between the estimated and the chronological age of the study participants, see Figure 1A).

**FIG. 1:**
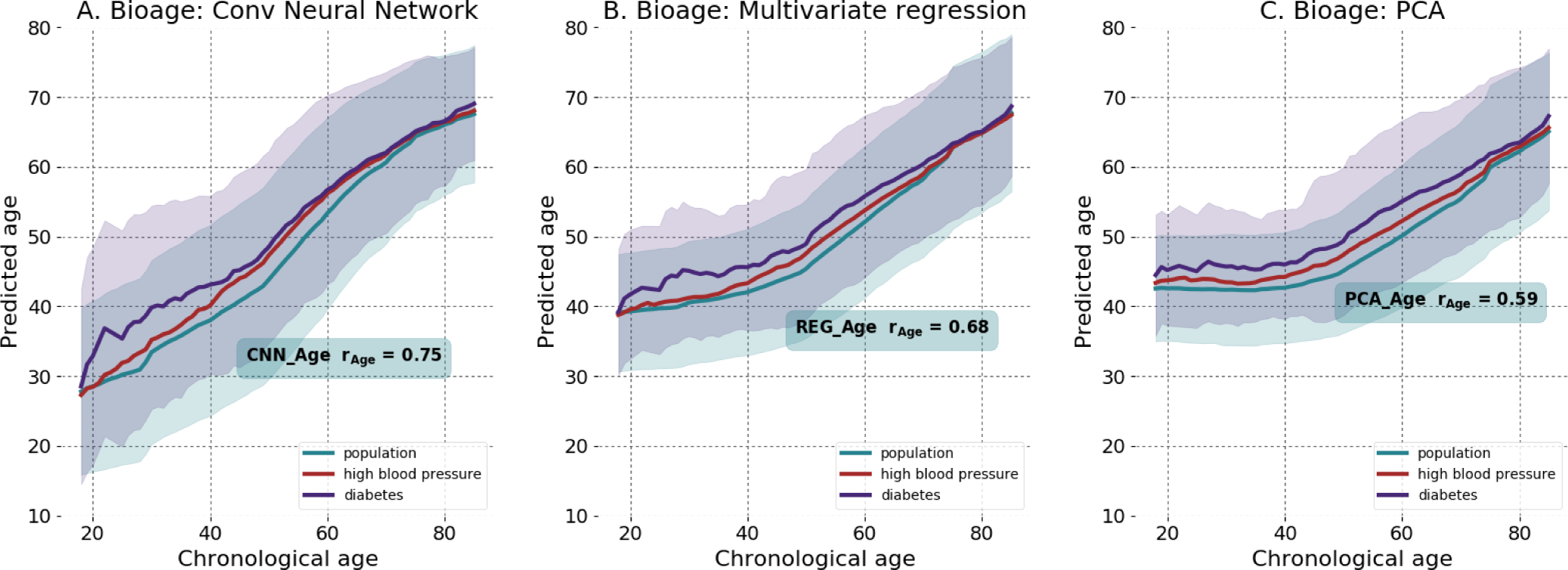
The biological age estimation according to deep convolution neural network CNN_Age (**A**), the multivariate regression REG_Age (**B**), and the unsupervised PCA_Age (**C**) models. The solid lines and the transparent standard deviation bands are color coded as green, blue, and red, representing the whole population, the patients diagnosed with diabetes by a doctor, and individuals with self-reported high blood pressure, respectively. All the calculations were produced using the NHANES 2003-2006 cohort wearable accelerometers data, comprising one-week long activity tracks (1*min^−1^* sampling rate).

To highlight the superior performance of the CNN_Age, we also characterized a regularized multivariate regression, trained to predict chronological age from a linear combination of hand-crafted features, representing statistical properties of the physical activities time series, borrowed from [5]. The result was the REG_Age model, which is similar by design to the most commonly used biological age metrics, such as, e.g., DNA methylation clock, a regularized linear regression of DNA methylation features to the chronological age [1, 18]. REG_Age worked fairly well, but was inferior in accuracy to CNN_Age (Pearson’s *r* = 0.68, see Figure 1B).

In our recent study [5] we observed that Principle Components (PC) analysis reveals that most of the variance in the multi-dimensional parameter space spanned by all NHANES participant representations (same as in REG_Age model) is associated with chronological age. Therefore, we proposed the first principal component score as the unsupervised definition of biological age, PCA_Age (Pearson’s correlation coefficient of the first PC score and chronological age *r* = 0.59, see Figure 1C). PCA_Age does not change much at first, but increases approximately linearly with age thereafter, roughly, the age of 40, and exhibits an excellent correlation with the negative logarithm of average daily activity [5].

CNN_Age models outperformed any other model in terms of the chronological age prediction accuracy. REG_Age and PCA_Age used the same set of physical activity derived features. Therefore, we suggest that the CNN produces the relevant set of age-associated properties of the raw time series in a fully automated way.

### Improved chronological age estimation accuracy undermines the biological age association with diseases and lifespan

Patients diagnosed with diabetes or hypertension appear to be biologically older than their healthy peers, see Figures 1A-C. To find out if the biological age difference translates into a lifespan change, we used NHANES death register representing survival data for 7837 participants (3750 male, 4087 female, aged 18 − 85, follow-up time up to 9 years, 701 participants died, in total). Figures 2A-C is a summary of Kaplan-Meier survival curves for NHANES participants, stratified into the highand the lowrisk groups according to the difference between the estimated biological age of an individual and the averaged estimated age of gender- and age-matched peers. The procedure is similar in spirit but formally different from and produces better results than the group separation according to the sign of aging acceleration. Using the predicted biological age as the reference naturally offsets the apparently non-linear relation between the biological and chronological age.

The unsupervised PCA_Age model fared exceptionally well and produced significantly different survival curves (*p* = 2 × 10^−7^, see Figure 2C). The increasing accuracy of chronological age estimation by each of the two supervised models, however, came at a price of a significant drop in the ability to distinguish the longerand the shorter-living individuals. The multivariate regression REG_Age and the deep CNN_Age models failed to rank the individuals according to mortality risks (*p* = 0.08 and *p* = 0.01, respectively, see Figures 2A and B).

**FIG. 2:**
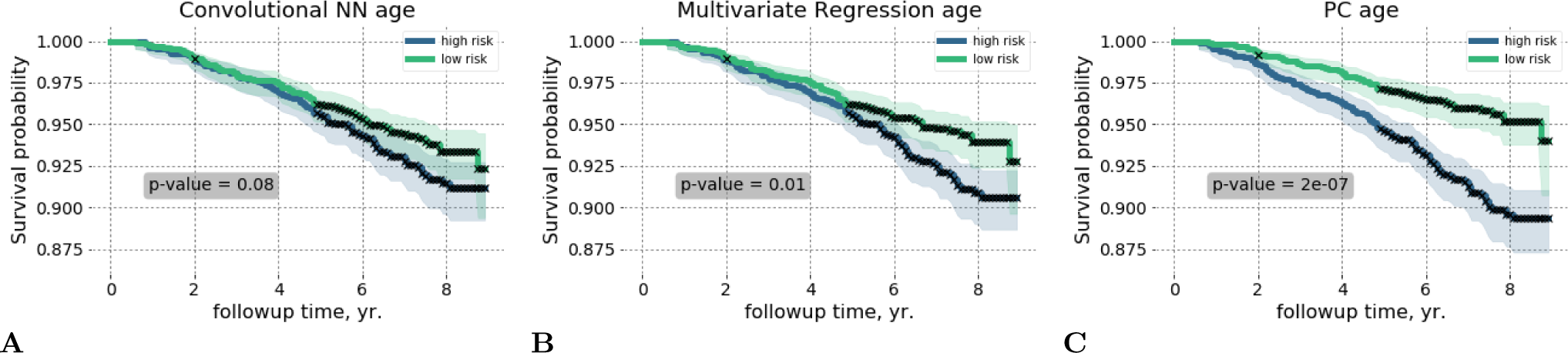
Kaplan-Meier survival curves illustrate qualitatively the performance of the biological age models to distinguish between the longerand the shorter-living individuals. Each participant was classified into the “high-” and the “low” risks groups according to the deep convolution neural network CNN_Age (**A**),the multivariate regression REG_Age (**B**), and the unsupervised PCA_Age (**C**) models. The p-values characterize the survival curves separation significance.

The apparent progressive loss of biologically relevant information by increasingly accurate models, involving a regression to chronological age, highlights the biological significance of aging acceleration, a quantity closely related to the chronological age determination error from the physiological data.

### Biological age contributes to health risk assessment

Health Risk Assessment (HRA) is a systematic approach to collecting information from individuals that identifies risk factors, provides individualized feedback, and links the person with health-promoting interventions, see, e.g., [19]. Biological age acceleration is associated with major diseases, and hence we asked ourselves if any of the biological age models could provide any useful additional information and improve health risks evaluation accuracy over standard HRA questionnaires.

HRA approaches involve capturing demographic (such as age and gender), lifestyle (including exercise, smoking, alcohol intake, diet), and physiological (e.g., weight, height, blood pressure, cholesterol) characteristics [19]. To model a HRA we built a Cox proportional hazards model including the most important demographic, lifestyle and physiological factors, see Table I for the summary of the model parameters. We then tested the statistical significance of the biological age estimators in association with all-cause mortality using a Cox proportional hazards model with the corresponding aging acceleration and the health risk factors taken as covariates, see the top of Table II for the summary of the results. The biological age scores were corrected by the health risks factors in advance to avoid possible fitting instabilities due to potential collinearity between the covariates. In this way, we explicitly tested the biological age residual for the ability to explain the risk function variance that is not already accounted for by the standard HRA factors.

**TABLE I:** Cox-Gompertz parametric proportional hazards model parameters.

| Covariate | Hazard Ratio, 95% CI, p-value |
| --- | --- |
| age | 1.090, CI[1.082, 1.099] ($p=2\text{e-}109$) |
| gender | 1.503, CI[1.281, 1.765] ( 2.4yr., $p=6\text{e-}07$) |
| diabetes | 1.690, CI[1.401, 2.038] ( 2.2yr., $p=4\text{e-}08$) |
| smoking | 2.300, CI[1.866, 2.836] ( 3.8yr., $p=6\text{e-}15$) |
| high blood pressure | 1.198, CI[1.018, 1.409] ( 1.0yr., $p=3\text{e-}02$) |
| log(activity) | 2.378, CI[2.051, 2.756] ( 4.9yr., $p=1\text{e-}30$) |

**TABLE II:**
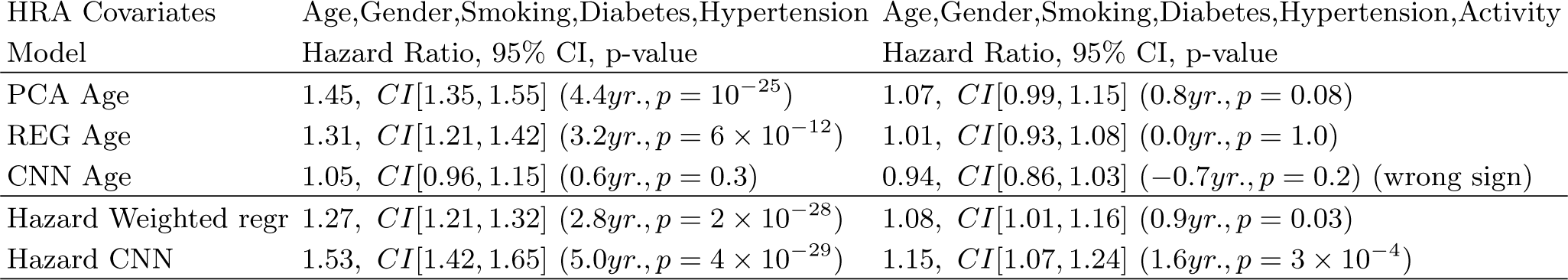
Association of the biological ageand all-causes mortality ‐predicting models with all-cause mortality. Two class of models were evaluated: one including the HRA parameters: age, gender, diabetes, smoking and hypertension as covariates (the left column) and the other including additionally the negative logarithm of the average daily physical activity (the biological age proxy, the right column). The corresponding 95% hazard ratio intervals are given along with the significance p-value and the effect levels, expressed in years of life lost (see the text for details).

| HRA Covariates Model | Age,Gender,Smoking,Diabetes,Hypertension Hazard Ratio, 95% CI, p-value | Age,Gender,Smoking,Diabetes,Hypertension,Activity Hazard Ratio, 95% CI, p-value |
| --- | --- | --- |
| PCA Age | 1.45, *CI*[1.35, 1.55] (4.4yr., $p = 10^{-25}$) | 1.07, *CI*[0.99, 1.15] (0.8yr., $p = 0.08$) |
| REG Age | 1.31, *CI*[1.21, 1.42] (3.2yr., $p = 6 \times 10^{-12}$) | 1.01, *CI*[0.93, 1.08] (0.0yr., $p = 1.0$) |
| CNN Age | 1.05, *CI*[0.96, 1.15] (0.6yr., $p = 0.3$) | 0.94, *CI*[0.86, 1.03] (-0.7yr., $p = 0.2$) (wrong sign) |
| Hazard Weighted regr | 1.27, *CI*[1.21, 1.32] (2.8yr., $p = 2 \times 10^{-28}$) | 1.08, *CI*[1.01, 1.16] (0.9yr., $p = 0.03$) |
| Hazard CNN | 1.53, *CI*[1.42, 1.65] (5.0yr., $p = 4 \times 10^{-29}$) | 1.15, *CI*[1.07, 1.24] (1.6yr., $p = 3 \times 10^{-4}$) |

As in every statistical test so far, the unsupervised PCA_Age model prediction produced the largest proportional hazards effect, *HR* = 1.45 (*p* < 10^−10^). The supervised REG_Age and CNN_Age models achieved better age predicting accuracy at the expense of biological significance loss, see the upper-left column in Table II. The multivariate regression REG_Age provided a smaller yet significant effect, HR = 1.31 (*p* < 10*^−10^*). The most accurate chronological age estimator CNN_Age, however, failed to produce any statistically significant contribution (*p* = 0.3). Therefore, at least in our model scenario, two of the biological age estimators, PCA_Age and REG_Age, produced biological age measures that substantially improve patient health risk estimation beyond any standard procedure involving HRA metrics.

On the other hand, the negative logarithm of the average daily physical activity is a good biological age proxy [5] and provides a highly significant contribution to the hazard function (*p* < 10*^−10^*), see Table I. We checked explicitly and found that none of the biological age models estimations produced any proportional hazards effect, once the physical activity level is included as an independent HRA variable, see the upper-right column of Table II, *p* > 0.05.

### Deep learning mortality model improves health risks assessment

Next, we explored whether we could build a useful health risks prediction model involving signatures of aging or diseases beyond biological age. Proportional hazards models (PHM) in Cox[20] or Cox-Gompertz[21] variants are trusted tools for such an analysis and are readily available as standard software, including a recent deep learning architecture implementation [22]. We employed CNN to automate the most relevant features extracted from the raw time series. Accordingly, we would need a PHM likelihood as the network cost function in a form, suitable for an efficient minimization with backpropagation. Naturally, we expected that the contribution to the all-cause mortality (or hazard ratio) of features, associated with the locomotor activity to be small on top of the already significant effects of age, gender, and the other HRA factors. Therefore, instead of implementing a full non-linear likelihood minimization, we performed a perturbation theory expansion of the Cox-Gompertz likelihood. The result was a simplified linear model, closely related to regression to Martingale residuals, or the unexplained risks variance of the model involving the standard HRA factors. More importantly, the cost function was formally equivalent to a linear regression with sample weights and therefore can be implemented out-of-the-box, with essentially any modern deep learning software (see Materials and Methods).

We used the proposed linearized version of the Cox-Gompertz proportional hazards model to train a deep CNN. The network received raw physical activity streams and performed considerably better than our reference, a linear hazard predictor built using the same perturbation theory expansion but with the help of hand-crafted features from [5] and already used here as descriptors by REG_Age and REG_Age models. The deep CNN hazard rate model outperformed all the biological age scores above, see the bottom of Table II. Both the linear and the CNN hazard rate predictors produced a significant association with the all-cause mortality after a correction by age, gender, diabetes, smoking and high blood pressure (*p* < 1*e* − 10, see the bottom-left column of Table II). The correction by the negative logarithm of the average daily physical activity made the hazard prediction of the linear hazard model statistically irrelevant. The CNN hazard rate model predictor remained significant, *HR* = 1.15, (*p* = 0.0003). Assuming proportional hazards model this contribution accounts for, approximately, 1.6 years of life gained or lost at the standard deviation level. The most significant association of the CNN hazard rate model residual after detrending by the physical activity level and the major HRA factors with the NHANES Questionnaire and Laboratory data variables was the self-reported “general health condition” (labels “excellent/very good” vs “fair/poor”).

## DISCUSSION

We report a systematic investigation of biological aging acceleration in relation to disease states and all-cause mortality in a large-scale human study. In particular, we used the NHANES physical activity records along with the medical meta-data and the death register to produce a series of biological age models. The most popular supervised learning examples, such as a multivariate regression and a deep CNN, exploit the apparent linear dependence of physiological changes with age and hence were trained to produce a chronological age estimate from individual locomotor activity time series. Both cases involved a minimization of the biological age acceleration, the difference between the “physiological” age, estimated by the model and the chronological age of a patient. Alternatively, we used an unsupervised technique taking advantage of the intrinsic low-dimensionality of the aging trajectories, closely related to criticality [5, 23] of the biological state variables kinetics, and thus presenting a natural biomarker of age from Principal Component Analysis.

To date, different forms of regularized multivariate linear regressions of biologically relevant variables to chronological age are the most popular technique behind the recently proposed biological age signatures, such as, e.g., biological clocks using IgG glycosylation [24], blood biochemical parameters [25], gut microbiota composition [26], and cerebrospinal fluid proteome [27]. The “epigenetic clock” based on DNA methylation (DNAm) levels [1, 18] appears to be the most accurate and the most extensively studied measure of aging. The biological age acceleration, measured by the DNAm clock, explains allcause mortality in later life better than chronological age [15], is elevated in people with HIV, Down syndrome [11, 12], obesity [13, 14], but is not correlated with smoking [28]. The same biomarker of age is lower for supercentennial’s offspring [16], and predicts mortality in a longitudinal twins study [17].

By design, such a supervised methodology involves a form of minimization of the biological age determination error, i.e., an attempt of minimizing the difference between the predicted “physiological” age and the chronological age of the patient. This is, by definition, the biological age acceleration, a presumably biologically relevant variable. Therefore, we expected and demonstrated here that a systematic improvement of the chronological age determination error minimization leads to an immediate degradation of the biological age acceleration significance in any test, involving health or risk of death. Our calculations show that the loss of biological information can be aggravated if even more powerful machine learning tools, such as a deep learning architecture, are employed to unravel complex and possibly nonlinear relations between the features in the data and produce even more “accurate” models. The most accurate chronological age estimation from biological samples could, however, find applications in forensic research [29].

Diabetes is one of the most significant health risk factors affecting lifespan by shortening life expectancy up to 8 years according to some studies [30]. However, the most popular DNAm clock did not label patients with diabetes diagnosed by a doctor as “biologically older” in at least one study [31]. In our studies, the patients diagnosed with diabetes and hypertension appear to be significantly older biological-age wise according to the unsupervised PCA_age model [5]. The multivariate regression REG_age is similar in spirit to DNAm age and produced a less significant separation between the groups of patients with the diseases and healthy controls. It remains to be seen, however, which of the interpretations would be supported with future versions of the DNAm clock, or, if our findings are specific to the source of the signal derived from human physical activity time series properties, known to be associated with both of the diseases.

Physical activity measurements recorded by wearable devices are, in theory, an ideal data source for building fully automated HRA systems for continuous health risk monitoring and real-time feedback to patients and care providers. The unsupervised PCA_age and a linear multivariate regression REG_age yielded valuable biological age models producing a biological aging acceleration estimate predictive of all-cause mortality, even after detrending by the HRA variables. The more accurate models, such as PCA_age and the deep CNN_age predictors explained a lesser degree of the death risk variation, respectively. We also observed, that an explicit addition in a HRA routine of a variable in high correlation with the biological age, such as the negative logarithm of the average daily activity, makes any of biological age predictors statistically irrelevant.

The idea of identifying patterns in biological signals in association with phenotypic differences is not new [32]. However, deep CNNs bring the idea to an entirely new level and are particularly useful for automation of the most relevant features engineering from the data in relation to human activity recognition (HAR) [33, 34], specific diseases or risks factors [10]. To demonstrate CNN capabilities for all-cause mortality evaluation from physical activity records, we introduced a novel way to train a proportional hazards model effectively. We observed that since the fundamental HRA factors such as age, gender, and major disease status already allow for the production of an excellent survival model. Therefore, the expected contribution of any combination of the locomotor activity derived features, after detrending by the HRA variables, would be expected to be small. Under the circumstances, the full mortality can be obtained by iterations, with the zero-order approximation model being the Cox-Gompertz mortality model, embracing the HRA descriptors as independent covariates. The first subsequent perturbation theory correction is then equivalent to a regression with sample weights, depending on the zero-order model parameters (a generalization of the method for a more generic non-parametric mortality model is also possible and will be reported elsewhere). The statistical power of the model is limited by the total number of death events, see Eq. (3) in Materials and Methods. In that view, ubiquitous deployment of wearable sensors promises unparalleled opportunities to achieve large population-wide coverage and thus to make possible the identification of additional smaller health risk effects at significance levels in the future.

The unsupervised PCA_Age yields an important insight on the dynamics of the physiological state in association with age. Biological age turns out to be related to an order-parameter, associated with the organism development and as such, undergoes a random walk on top of the systematic aging drift [5, 23]. The variance of the biological age distribution increases with age, which is a sign of the increasing heterogeneity of the human population. The effect is a challenge to supervised methods, such as regressions to biological age and even proportional hazards models. We envision future improvements of mortality prediction models by taking into account the diffusion of the biological variables into the likelihood function directly. Given the success of the unsupervised biological age model in our study, we further expect a development of unsupervised deep learning architectures, such as deep auto-encoders [35] for aging research.

Life and health insurance programs have begun to provide discounts to their users based on physical activity monitored by fitness wristbands [36]. We report that a deep CNN can be used to further refine the risks models by inclusion of an apparently biological age-independent risk factor, producing a significant effect on lifespan. We believe that the result highlights the power and practical utility of semi-analytic approaches, combining aging theory with the most powerful modern machine learning tools. This synthesis will eventually produce even better health risks models for HRA, to mitigate longevity risks in insurance, help in pension planning, and contribute to upcoming clinical trials and future deployment of antiaging therapies.

## CONCLUSIONS

We performed a systematic evaluation of biological age models built from the data, representing physical activity tracks from a large cross-section human study. Our findings support the biological relevance of aging acceleration, the quantity often minimized in supervised biological models. We show that an apparent model accuracy improvement may come at a price of substantial degradation of a biological age utility in all applications involving health or all-cause mortality risks.

We propose a simple and efficient way to train parametric proportional hazards models using out-of-the-box deep machine learning software. We trained and characterized a proof-of-concept deep CNN hazard rate model to demonstrate fully automated feature engineering from complex biological signals analysis and produce biomarkers of age and mortality at the level of accuracy exceeding that of the traditional approaches.

## MATERIALS AND METHODS

### Data preparation, quantification of locomotor activity

Locomotor activity records and questionnaire/laboratory data from the National Health and Nutrition Examination Survey (NHANES) 2003-2004 and 2005-2006 cohorts were downloaded from [www.cdc.gov/nchs/nhanes/index.htm]. NHANES provides locomotor activity in the form of 7-day long continuous tracks of “activity counts” sampled at 1*min^−1^* frequency and recorded by a physical activity monitor (ActiGraph AM-7164 single-axis piezoelectric accelerometer) worn on the hip. Of 14,631 study participants (7176 in the 2003-2004 cohort and 7455 in the 2005-2006 cohort), we filtered out samples with abnormally low (average activity count <50) or high (>5000) physical activity. We also excluded participants aged 85 and older since the NHANES age data field is top coded at 85 years of age and we desired precise age information for our study. The mortality data for NHANES participants is obtained from the National Center for Health Statistics public resources (4017 in the 2003-2004 cohort and 3985 in the 2005-2006 cohort). We excluded days with less than 200 minutes corresponding to activity states > 0. Only participants with 4 or more days that passed this additional filter were retained, yielding a total of 7454 samples (age, years: 35 ± 23, range 6 − 84; women: 51%). Quantification of locomotor activity of the subject was carried out in two ways: by hand-engineered features for all linear analysis and by CNN (see corresponding sections below).

### Statistical description time series representing physical activity

To calculate a statistical descriptor of each participant’s locomotor activity we followed the prescription from [5]. We first converted activity counts into discrete states with bin edges *b_k_*, *k = 1…K*. Activity level states 1*…K* − 1 were then defined as half-open intervals *b_k_ ≤ a* < *b*_*k*+1_, state 0 as *a < b*_1_ and state *K* as *a ≥ b_K_*, where *a* is the activity count value. In this study, we defined *K* = 8 activity states with bin edges *b_k_* = *e^k^* 1., k = 1….7. Thus, each sample was converted into a track of activity states and a transition matrix (TM) was then calculated for each participant (see below). To ensure that our analysis dealt only with days on which a participant actually performed some physical activity, we applied an additional filter. Transition matrices (TM) *T_ij_, i* = 1…8*, j* = 1…8 were calculated as a set of transition rates from each state *j* to each other state *i* (the diagonal elements correspond to the probability of remaining in the same activity state). We flattened 8 × 8 TM of each sample into a 64-dimensional descriptor vector and converted the flattened descriptor to log-scale to ensure approximately normal distribution for elements of the locomotor descriptor (a useful property for the stability of the linear models that we applied in PCA and Survival analysis). All near-zero elements (< 10^-3^, which corresponds to less than 10 transitions during a week) were imputed by the value of 10^-3^ before log-scaling.

### Age predicting models

We compare three age-estimating models. CNN_Age is a convolution neural network (CNN) model trained to predict the age of an individual based on input raw activity counts track (the details of CNN architecture are described below).

The remaining models, REG_Age and PCA_Age, estimate age in response to the vector of hand-crafted features representing statistical properties of the physical activity time series and borrowed from [5]. We trained the models using the activity records of 7186 NHANES participants aged 18 – 85. REG_Age is a *l*_2_ regularized multivariate regression trained with 5-fold cross-validation using RidgeCV package from scikit-learn python library.

PCA_Age is based on principal component (PC) analysis of the same set of features with the help of SVD factorization from numpy python library. We retained the first principal component score as the biological age metric and reported either the raw PC score or a variable scaled to age by a univariate linear regression.

### Cox-Gompertz proportional hazards model

According to the Gompertz law [37], the mortality rate in human populations increases exponentially starting at the age of about 40. Therefore an accurate estimation of a hazard rate, or mortality, for each participant can be obtained with the help of a parametric Cox-Gompertz proportional hazards model adapted from [21]. The model predicts the mortality rate in the form *M* (*t^n^, x^n^*) = *M*_0_ exp(Γ*t^n^*) exp(*β, x^n^*), where *t^n^* is the age of a participant *n*, *x^n^* is a vector of independent predictor variables (covariates), such as the participant’s gender and any set of physical activity related descriptors. The variables Γ and *M*_0_ stand for the Gompertz exponent (inversely related to the mortality rate doubling time) and the initial mortality rate, respectively. The unknown parameters along with the vector of quantities *β* are to be fitted using the experimental data by minimizing the negative of the following log-likelihood function:

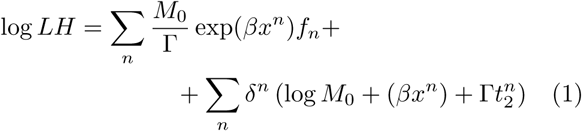

where the summation occurs over all the study participants, 
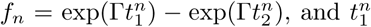
 is the age of a participant when locomotor activity measurements were carried out. The second time variables, 
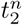
, is the age at death, if the patient died during the follo-up time or the age of the last observation if the individual was alive, respectively. The indicator *δ^n =^* 1 if a patient *n* is dead at 
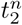
 and *δ^n =^* 0 otherwise. The log-likelihood can be further regularized by adding a proper term depending on a norm of the vector *β* (*L*_2_*−* regularization in our study). The optimization with respect to the scalar variables Γ, *M*_0_ and the vector *β* can be performed using any convenient technique, including Broyden-Fletcher-Goldfarb-Shanno (BFGS) algorithm or a batched stochastic gradient descent (SGD).

## Perturbation theory expansion in a Cox-Gompertz model

We start assuming there’s a well-defined minimum defined by a set of the best fit variables Γ, *M*_0_ and *β*¯. Let us now see how the model can be improved if we are allowed to add a set of additional variables *ξ* to improve the hazard rate model. Once the new parameters are plugged in the likelihood function (1), then, the position of the minimum of the likelihood function (1) also changes. If the corrections to the model parameters *δ*Γ, *δM*_0_, *δβ* are sufficiently small, the solution of the minimization problem can be obtained by iterations. In typically small cohorts of human subjects, the modifications of the Gompertzian variables *δ*Γ and *δM*_0_ are poorly constrained [38]. Therefore the variation of each one of the parameters can be arbitrary set to zero. We choose to fix Γ and without further derivation provide the final expression for the proportional hazards effect:

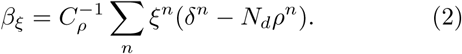

Here 
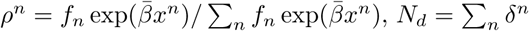
 is the total number of the death events in the dataset, and 
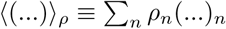
 stands for averaging with the sample weights *ρ^n^*. By definition the sample-weighted variance 
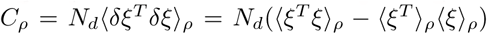
. The effect estimation error can also be obtained analytically,

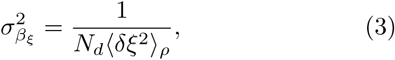

and depends explicitly on the number of death events, rather than on the total number of individuals, in the study.

Up to the appearance of sample-weighted average in the definition of the covariance matrix, Eq. (2) is a regression of independent variables *ξ* to the martingale residual of the Cox-Gompertz model. The latter is the difference between the actual survival, represented by the indicator variable *δ^n^*, and the model mortality, integrated over the duration of the follow-up time for the same patient, 
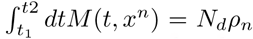
. What is more important, the solution can now be obtained by minimization of an extremely simple cost function

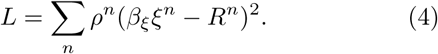

This is nothing else but a sample-weighted regression of *ξ^n^* against the properly selected target function *R^n^* = 1 − *δ^n^*/(*ρ^n^N_d_*). This form of the likelihood optimization can be easily done with any modern software library and hence is far more convenient for machine learning applications than the original Cox-Gompertz likelihood (1).

## Significance of locomotor hazards and bioage in all-cause mortality evaluation

We first tested the age and hazard predicting models for the significance of their association with allcause mortality with the help of a Cox proportional hazards model. Every test included the model prediction for each NHANES participant along with the age (data field RIDAGEMN), gender (RIAGENDR), diabetes (DIQ010), smoking (SMQ040), and hypertension (high blood pressure) (BPQ020). Also, the same test was carried out after including two additional covariates in the form of the average number of activity counts per day and the logarithm of this number, calculated from the physical activity recording tracks. Each covariate except for age were subsequently linearly detrended by all the other covariates and then standardized to zero mean and unit variance. After these preprocessing procedures, the significance for association was tested using Cox proportional hazards model yielding the effect, its 95% confidence intervals (CI) and p-value. The effect was further transformed into the corresponding difference in life expectancy by dividing the effect by the Gompertz exponential coefficient 0.085 *yrs^−1.^*.

The Cox model parameters and the significance pvalues are summarized in Table II and were obtained using *survival* [39, 40] package implemented in *R* [41]. Alternatively, we obtained the same p-values as probabilities of the effect deviations from zero using the analytically expressions for the effect and the effect determination error given by Eqs. (2) and (3).

## Convolution neural networks for biological age and risks of death predictions

We trained a convolution neural network to convert each physical activity record, a vector of 10080 values, representing subsequent activity counts per minute records (7-day-long recording sampled at 1*min^−1^* frequency), into a single value of estimated age or hazard rate. The network architecture was similar for both tasks and is shown in Figure 3. We identified an optimal architecture for this deep learning model empirically, after exploring a range of combinations of layer depth and size. Before feeding data into convolution network, we apply Batch Normalization [42]. The network consisted of four convolution layers with ReLU activation each followed by Max Pooling. The results were imported into two fully connected layers with ReLU activation. We applied the dropout [43] to regularize the dense layers. We used the RmsProp [44] optimizer with learning rate per sample set to 10*^−7^*, minimal learning rate 10*^−17^* and momentum 0.9. Dropout rate is set to 0.5 after first 9 epochs, and no dropout was used during these 9 epochs. Finally, output of the last layer is fed into a single linear neuron (a linear regression to the network target) producing the resulting value of age prediction.

**FIG. 3:**
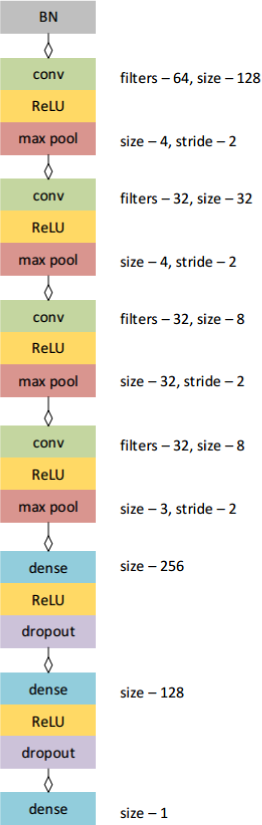
Architecture of convolution neural network.

The convolution layers and output dense layer weights were initialized with Glorot uniform initializer [45]. We used a gaussian initializer for the two internal dense layers weights. For CNN_Age model we scaled the weights of the output dense layer by the factor 85 to match the range of the possible output values.

To produce a deep CNN for mortality prediction using the linearity log-likelihood (4), we started by building the simplest “zero-order” Cox-Gompertz PHM including the participant’s age and gender as independent covariate. In spite of the very small number of death cases in the data, the Gompertz exponent Γ = 0.088 was compatible with the currently accepted mortality rate Γ = 0.085, corresponding to doubling time of 8 years [46]. The initial mortality rate was *M*_0_ = 3 × 10^−5^ *yrs^−1.^*, and therefore the model predicted 90.02 years of life expectancy, on average (gender contributed 5.2 years of the life expectancy difference).

A CNN trained to predict the linearized proportional hazards model target *R^n^* from the raw physical activity tracks generated the best locomotor activity features *ξ* (output of the last hidden CNN layer) and identified the projection vector *β_ξ_* (weights of last linear layer). The values *R^n^* were inverse normal transformed and standardized to zero mean and unit variance with custom *R* [41] script. We tested the CNN output for significance by feeding it to a Cox model along with age, gender, and major risk factors: smoking, diabetes, high blood pressure and, optionally, average daily activity. We observed that a progressive training of the CNN reaches a plateau in significance for mortality prediction starting from training epoch m2500 (data not shown).

## ACKNOWLEDGEMENTS

The work has been funded by Gero LLC. The authors thank their colleagues from Gero K. Avchaciov, A. Tarkhov, D. Shishov, and G. Getmantsev for extensive help with handling the data and proof-reading of the work.

## References

[1] G. Hannum, J. Guinney, L. Zhao, L. Zhang, G. Hughes, S. Sadda, B. Klotzle, M. Bibikova, J. B. Fan, Y. Gao, R. Deconde, M. Chen, I. Rajapakse, S. Friend, T. Ideker, and K. Zhang, “Genome-wide methylation profiles reveal quantitative views of human aging rates,” Mol. Cell, vol. 49, pp. 359–367, Jan 2013.

[2] M. J. Peters, R. Joehanes, L. C. Pilling, C. Schurmann, K. N. Conneely, J. Powell, E. Reinmaa, G. L. Sutphin, A. Zhernakova, K. Schramm, Y. A. Wilson, S. Kobes, T. Tukiainen, Y. F. Ramos, H. H. Goring, M. Fornage, Y. Liu, S. A. Gharib, B. E. Stranger, P. L. De Jager, A. Aviv, D. Levy, J. M. Murabito, P. J. Munson, T. Huan, A. Hofman, A. G. Uitterlinden, F. Rivadeneira, J. van Rooij, L. Stolk, L. Broer, M. M. Verbiest, M. Jhamai, P. Arp, A. Metspalu, L. Tserel, L. Milani, N. J. Samani, P. Peterson, S. Kasela, V. Codd, A. Peters, C. K. Ward-Caviness, C. Herder, M. Waldenberger, M. Roden, P. Singmann, S. Zeilinger, T. Illig, G. Homuth, H. J. Grabe, H. Volzke, L. Steil, T. Kocher, A. Murray, D. Melzer, H. Yaghootkar, S. Bandinelli, E. K. Moses, J. W. Kent, J. E. Curran, M. P. Johnson, S. Williams-Blangero, H. J. Westra, A. F. McRae, J. A. Smith, S. L. Kardia, I. Hovatta, M. Perola, S. Ripatti, V. Salomaa, A. K. Henders, N. G. Martin, A. K. Smith, D. Mehta, E. B. Binder, K. M. Nylocks, E. M. Kennedy, T. Klengel, J. Ding, A. M. Suchy-Dicey, D. A. Enquobahrie, J. Brody, J. I. Rotter, Y. D. Chen, J. Houwing-Duistermaat, M. Kloppenburg, P. E. Slagboom, Q. Helmer, W. den Hollander, S. Bean, T. Raj, N. Bakhshi, Q. P. Wang, L. J. Oyston, B. M. Psaty, R. P. Tracy, G. W. Montgomery, S. T. Turner, J. Blangero, I. Meulenbelt, K. J. Ressler, J. Yang, L. Franke, J. Kettunen, P. M. Visscher, G. G. Neely, R. Korstanje, R. L. Hanson, H. Prokisch, L. Ferrucci, T. Esko, A. Teumer, J. B. van Meurs, A. D. Johnson, M. A. Nalls, D. G. Hernandez, M. R. Cookson, R. J. Gibbs, J. Hardy, A. Ramasamy, A. B. Zonderman, A. Dillman, B. Traynor, C. Smith, D. L. Longo, D. Trabzuni, J. Troncoso, M. van der Brug, M. E. Weale, R. OBrien, R. Johnson, R. Walker, R. H. Zielke, S. Arepalli, M. Ryten, and A. B. Singleton, “The transcriptional landscape of age in human peripheral blood,” Nat Commun, vol. 6, p. 8570, Oct 2015.

[3] S. Enroth, S. B. Enroth, A. Johansson, and U. Gyllensten, “Protein profiling reveals consequences of lifestyle choices on predicted biological aging.,” Sci Rep, vol. 5, p. 17282, Dec. 2015.

[4] B. C. Choi, A. W. Pak, and J. C. Choi, “Daily step goal of 10,000 steps: a literature review,” Clinical & Investigative Medicine, vol. 30, no. 3, pp. 146–151, 2007.

[5] T. V. Pyrkov, E. Getmantsev, B. Zhurov, K. Avchaciov, M. Pyatnitskiy, L. Menshikov, K. Khodova, A. V. Gudkov, and P. O. Fedichev, “Quantitative characterization of biological age and frailty based on locomotor activity records,” bioRxiv, p. 186569, 2017.

[6] P. Rajpurkar, A. Y. Hannun, M. Haghpanahi, C. Bourn, and A. Y. Ng, “Cardiologist-level arrhythmia detection with convolutional neural networks,” arXiv preprint arXiv:1707.01836, 2017.

[7] E. Putin, P. Mamoshina, A. Aliper, M. Korzinkin, A. Moskalev, A. Kolosov, A. Ostrovskiy, C. Cantor, J. Vijg, and A. Zhavoronkov, “Deep biomarkers of human aging: application of deep neural networks to biomarker development,” Aging (Albany NY), vol. 8, no. 5, p. 1021, 2016.

[8] A. A. Cohen, V. Morissette-Thomas, L. Ferrucci, and L. P. Fried, “Deep biomarkers of aging are population-dependent,” Aging (Albany NY), vol. 8, no. 9, p. 2253, 2016.

[9] Z. Wang, L. Li, B. S. Glicksberg, A. Israel, J. T. Dudley, and A. Ma’ayan, “Predicting age by mining electronic medical records with deep learning characterizes differences between chronological and physiological age,” Journal of Biomedical Informatics, 2017.

[10] L. Oakden-Rayner, G. Carneiro, T. Bessen, J. C. Nascimento, A. P. Bradley, and L. J. Palmer, “Precision radiology: Predicting longevity using feature engineering and deep learning methods in a radiomics framework,” Scientific Reports, vol. 7, no. 1, p. 1648, 2017.

[11] S. Horvath and A. J. Levine, “HIV-1 Infection Accelerates Age According to the Epigenetic Clock.,” J. Infect. Dis., vol. 212, pp. 1563–73, Nov. 2015.

[12] S. Horvath, P. Garagnani, M. G. Bacalini, C. Pirazzini, S. Salvioli, D. Gentilini, A. M. D. Blasio, C. Giuliani, S. Tung, H. V. Vinters, and C. Franceschi, “Accelerated epigenetic aging in Down syndrome.,” Aging Cell, vol. 14, pp. 491–5, June 2015.

[13] S. Horvath, W. Erhart, M. Brosch, O. Ammerpohl, W. von Schönfels, M. Ahrens, N. Heits, J. T. Bell, P.-C. Tsai, T. D. Spector, et al., “Obesity accelerates epigenetic aging of human liver,” Proceedings of the National Academy of Sciences, vol. 111, no. 43, pp. 15538–15543, 2014.

[14] S. Horvath, W. Erhart, M. Brosch, O. Ammerpohl, W. von SchÃűnfels, M. Ahrens, N. Heits, J. T. Bell, P. C. Tsai, T. D. Spector, P. Deloukas, R. Siebert, B. Sipos, T. Becker, C. RÃűcken, C. Schafmayer, and J. Hampe, “Obesity accelerates epigenetic aging of human liver.,” Proc. Natl. Acad. Sci. U.S.A., vol. 111, pp. 15538–43, Oct. 2014.

[15] R. E. Marioni, S. Shah, A. F. McRae, B. H. Chen, E. Colicino, S. E. Harris, J. Gibson, A. K. Henders, P. Redmond, S. R. Cox, et al., “Dna methylation age of blood predicts all-cause mortality in later life,” Genome biology, vol. 16, no. 1, p. 25, 2015.

[16] S. Horvath, C. Pirazzini, M. G. Bacalini, D. Gentilini, A. M. D. Blasio, M. Delledonne, D. Mari, B. Arosio, D. Monti, G. Passarino, F. D. Rango, P. D’Aquila, C. Giuliani, E. Marasco, S. Collino, P. Descombes, P. Garagnani, and C. Franceschi, “Decreased epigenetic age of PBMCs from Italian semi-supercentenarians and their offspring.,” Aging (Albany NY), vol. 7, pp. 1159–70, Dec. 2015.

[17] L. Christiansen, A. Lenart, Q. Tan, J. W. Vaupel, A. Aviv, M. McGue, and K. Christensen, “DNA methylation age is associated with mortality in a longitudinal Danish twin study.,” Aging Cell, vol. 15, pp. 149–54, Feb. 2016.

[18] S. Horvath, “DNA methylation age of human tissues and cell types,” Genome Biol., vol. 14, no. 10, p. R115, 2013.

[19] J. M. Stellman, Encyclopaedia of occupational health and safety. International Labour Organization, 1998.

[20] D. R. Cox, “Regression models and life-tables,” in Breakthroughs in statistics, pp. 527–541, Springer, 1992.

[21] B. Efron, “The efficiency of cox’s likelihood function for censored data,” Journal of the American statistical Association, vol. 72, no. 359, pp. 557–565, 1977.

[22] J. Katzman, U. Shaham, J. Bates, A. Cloninger, T. Jiang, and Y. Kluger, “Deep survival: A deep cox proportional hazards network,” arXiv preprint arXiv:1606.00931, 2016.

[23] D. Podolskiy, I. Molodtcov, A. Zenin, V. Kogan, L. Menshikov, R. J. S. Reis, and P. Fedichev, “Critical dynamics of gene networks is a mechanism behind ageing and gompertz law,” arXiv preprint arXiv:1502.04307, 2015.

[24] J. KriÅątiÄĞ, F. VuÄŊkoviÄĞ, C. Menni, L. KlariÄĞ, T. Keser, I. Beceheli, M. PuÄŊiÄĞ-BakoviÄĞ, M. Novokmet, M. Mangino, K. Thaqi, P. Rudan, N. Novokmet, J. Sarac, S. Missoni, I. KolÄŊiÄĞ, O. PolaÅąek, I. Rudan, H. Campbell, C. Hayward, Y. Aulchenko, A. Valdes, J. F. Wilson, O. Gornik, D. Primorac, V. ZoldoÅą, T. Spector, and G. Lauc, “Glycans are a novel biomarker of chronological and biological ages.,” J. Gerontol. A Biol. Sci. Med. Sci., vol. 69, pp. 779–89, July 2014.

[25] M. E. Levine, “Modeling the rate of senescence: can estimated biological age predict mortality more accurately than chronological age?,” J. Gerontol. A Biol. Sci. Med. Sci., vol. 68, pp. 667–674, Jun 2013.

[26] T. Odamaki, K. Kato, H. Sugahara, N. Hashikura, S. Takahashi, J. Z. Xiao, F. Abe, and R. Osawa, “Agerelated changes in gut microbiota composition from newborn to centenarian: a cross-sectional study,” BMC Microbiol., vol. 16, p. 90, May 2016.

[27] G. S. Baird, S. K. Nelson, T. R. Keeney, A. Stewart, S. Williams, S. Kraemer, E. R. Peskind, and T. J. Montine, “Age-dependent changes in the cerebrospinal fluid proteome by slow off-rate modified aptamer array.,” Am. J. Pathol., vol. 180, pp. 446–56, Feb. 2012.

[28] X. Gao, Y. Zhang, K.-U. Saum, B. Schöttker, L. P. Breitling, and H. Brenner, “Tobacco smoking and smoking related dna methylation are associated with the development of frailty among older adults,” Epigenetics, no. just accepted, 2016.

[29] A. Vidaki, D. Ballard, A. Aliferi, T. H. Miller, L. P. Barron, and D. S. Court, “Dna methylation-based forensic age prediction using artificial neural networks and next generation sequencing,” Forensic Science International: Genetics, vol. 28, pp. 225–236, 2017.

[30] O. H. Franco, E. W. Steyerberg, F. B. Hu, J. Mackenbach, and W. Nusselder, “Associations of diabetes mellitus with total life expectancy and life expectancy with and without cardiovascular disease,” Archives of internal medicine, vol. 167, no. 11, pp. 1145–1151, 2007.

[31] S. Horvath, M. Gurven, M. E. Levine, B. C. Trumble, H. Kaplan, H. Allayee, B. R. Ritz, B. Chen, A. T. Lu, T. M. Rickabaugh, et al., “An epigenetic clock analysis of race/ethnicity, sex, and coronary heart disease,” Genome biology, vol. 17, no. 1, p. 171, 2016.

[32] A. E. Brown, E. I. Yemini, L. J. Grundy, T. Jucikas, and W. R. Schafer, “A dictionary of behavioral motifs reveals clusters of genes affecting caenorhabditis elegans locomotion,” Proceedings of the National Academy of Sciences, vol. 110, no. 2, pp. 791–796, 2013.

[33] F. J. Ordóñez and D. Roggen, “Deep convolutional and lstm recurrent neural networks for multimodal wearable activity recognition,” Sensors, vol. 16, no. 1, p. 115, 2016.

[34] Y. Guan and T. Ploetz, “Ensembles of deep lstm learners for activity recognition using wearables,” arXiv preprint arXiv:1703.09370, 2017.

[35] G. E. Hinton and R. R. Salakhutdinov, “Reducing the dimensionality of data with neural networks,” science, vol. 313, no. 5786, pp. 504–507, 2006.

[36] S. Tedesco, J. Barton, and B. OâĂŹFlynn, “A review of activity trackers for senior citizens: Research perspectives, commercial landscape and the role of the insurance industry,” Sensors, vol. 17, no. 6, p. 1277, 2017.

[37] B. Gompertz, “On the nature of the function expressive of the law of human mortality, and on a new mode of determining the value of life contingencies,” Philosophical transactions of the Royal Society of London, vol. 115, pp. 513–583, 1825.

[38] A. E. Tarkhov, L. I. Menshikov, and P. O. Fedichev, “Strehler-mildvan correlation is a degenerate manifold of gompertz fit,” Journal of theoretical biology, vol. 416, pp. 180–189, 2017.

[39] T. M. Therneau, A Package for Survival Analysis in S, 2015. version 2.38.

[40] Terry M. Therneau and Patricia M. Grambsch, Modeling Survival Data: Extending the Cox Model. New York: Springer, 2000.

[41] R Core Team, R: A Language and Environment for Statistical Computing. R Foundation for Statistical Computing, Vienna, Austria, 2017.

[42] S. Ioffe and C. Szegedy, “Batch normalization: Accelerating deep network training by reducing internal covariate shift,” in International Conference on Machine Learning, pp. 448–456, 2015.

[43] G. E. Hinton, N. Srivastava, A. Krizhevsky, I. Sutskever, and R. R. Salakhutdinov, “Improving neural networks by preventing co-adaptation of feature detectors,” arXiv preprint arXiv:1207.0580, 2012.

[44] T. Tieleman and G. Hinton, “Rmsprop: Divide the gradient by a running average of its recent magnitude. coursera: Neural networks for machine learning,” tech. rep., Technical report, 2012. 31.

[45] X. Glorot and Y. Bengio, “Understanding the difficulty of training deep feedforward neural networks,” in Proceedings of the Thirteenth International Conference on Artificial Intelligence and Statistics, pp. 249–256, 2010.

[46] L. B. Levy G, The Biostatistics of Aging: From Gompertzian Mortality to an Index of Aging-relatedness. John Wiley & Sons, 2014.

